# Revisiting the phylogeny of Zoanthidea (Cnidaria: Anthozoa): staggered alignment of hypervariable sequences improves species tree inference

**DOI:** 10.1101/161117

**Authors:** Timothy D. Swain

## Abstract

The recent rapid proliferation of novel taxon identification in the Zoanthidea has been accompanied by a parallel propagation of gene trees as a tool of species discovery, but not a corresponding increase in our understanding of phylogeny. This disparity is caused by the trade-off between the capabilities of automated DNA sequence alignment and data content of genes applied to phylogenetic inference in this group. Conserved genes or segments are easily aligned across the order, but produce poorly resolved trees; hypervariable genes or segments contain the evolutionary signal necessary for resolution and robust support, but sequence alignment is daunting. Staggered alignments are a form of phylogeny-informed sequence alignment composed of a mosaic of local and universal regions that allow phylogenetic inference to be applied to all nucleotides from both hypervariable and conserved gene segments. Comparisons between species tree phylogenies inferred from all data (staggered alignment) and hypervariable-excluded data (standard alignment) demonstrate improved confidence and greater topological agreement with other sources of data for the complete-data tree. This novel phylogeny is the most comprehensive to date (in terms of taxa and data) and can serve as an expandable tool for evolutionary hypothesis testing in the Zoanthidea.

**Resumen:** Spanish language translation by Lisbeth O. Swain, DePaul University, Chicago, Illinois, 60604, USA.

Aunque la proliferación reciente y acelerada en la identificación de taxones en Zoanthidea ha sido acompañada por una propagación paralela de los árboles de genes como una herramienta en el descubrimiento de especies, no hay una correspondencia en cuanto a la ampliación de nuestro conocimiento en filogenia. Esta disparidad, es causada por la competencia entre la capacidad de los alineamientos de secuencia del ácido desoxirribonucleico (ADN) automatizados y la información contenida en los datos de genes que se aplican a los métodos de inferencia filogenética en este grupo de Zoanthidea. Las regiones o segmentos de genes conservados son fácilmente alineados dentro del orden; sin embargo, producen árboles de genes con resultados paupérrimos; además, aunque estas regiones hipervariables de genes o segmentos contienen las señas evolutivas necesarias para apoyar la construcción robusta y completa de árboles filogenéticos, estos genes producen alineamientos de secuencia abrumadores. Los alineamientos escalonados de secuencias son una forma de alineamientos informados por la filogenia y compuestos de un mosaico de regiones locales y universales que permiten que inferencias filogenéticas sean aplicadas a todos los nucleótidos de regiones hipervariables y de genes o segmentos conservados. Las comparaciones entre especies de árboles filogenéticos que se infirieron de los datos de alineamientos escalonados y los datos hipervariables excluidos (alineamiento estandarizado), demuestran un mejoramiento en la confiabilidad y un mayor acuerdo tipológico con respecto a otras fuentes que contienen árboles filogenéticos hechos de datos más completos. Esta nueva forma escalonada de filogenia es una de los más compresibles hasta la fecha (en términos de taxones y datos) y que pueden servir como una herramienta de amplificación para probar la hipótesis evolutiva de Zoanthidea.

## 1. Introduction

Nucleotide sequence-based molecular phylogenetics have been intensively applied to the Anthozoa order Zoanthidea, however our understanding of the evolutionary relationships among zoanthidean species has not progressed at the same pace. Since 2004, at least 107 phylogenetic trees that focus mostly or exclusively on Zoanthidea have been published in 46 reports (a rate of nearly 9 trees and 4 reports per year; Table 1). Within these trees, the mean number of species per tree is only 16.2, while the mean number of terminals is nearly three times as large, with some published trees containing <5% unique species among their terminals (Table 1). Furthermore, of the 107 trees identified, 100 (or 94%) are gene trees relying upon a single gene or gene fragment rather than attempted species trees (see section 3.1), and 30% of these gene trees are built solely on data from the mitochondrial cytochrome oxidase subunit I gene (COI; Table 1). It has been known since at least 2002 that the rate of evolution of COI within Anthozoa is insufficient for distinguishing between closely related species and is largely uninformative for addressing phylogenetic questions below the family or genus-level (Shearer et al., 2002). The details of these published trees suggest that the primary goal of phylogenetic research on this order could not be the relationships among species, but is more likely the exploration of gene evolution among specimens (see section 3.1). However, these gene trees have been generally used in species delimitation and are subsequently over-interpreted as species trees and molecular parataxonomic evidence of evolutionary relationships among species and higher taxa in the nearly complete absence of other data (see Swain et al., 2015; Swain et al., 2016; Swain and Swain, 2014 for further discussion of molecular parataxonomy).

**Table 1.**
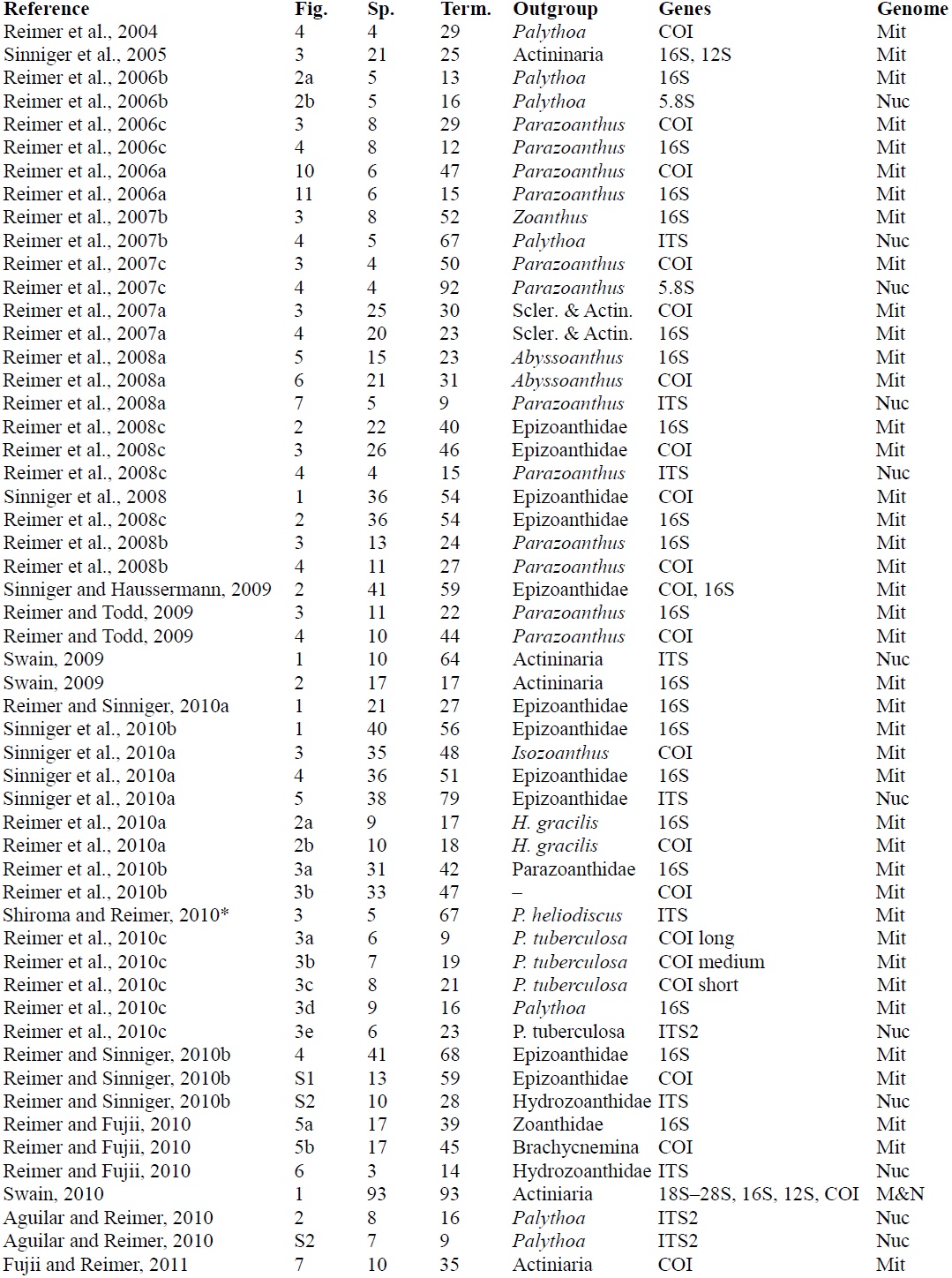
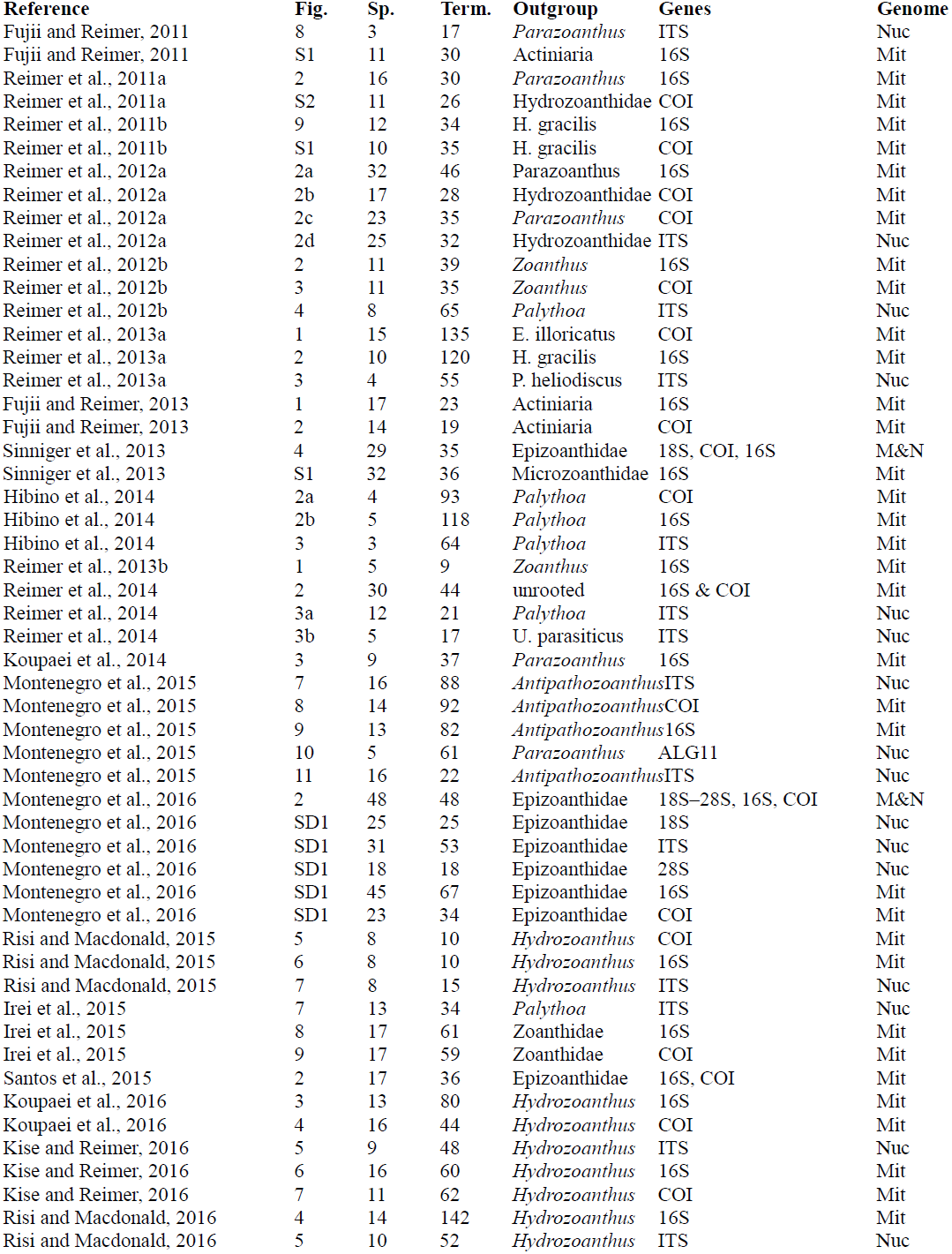
Review of published Zoanthidea molecular phylogenies inferred from nucleotide sequences, including the source reference (Reference), figure number of the phylogeny within that source (Fig.), number of apparent species in each phylogeny (Sp.), number of terminals in each phylogeny (Term.), outgroup employed in each phylogeny (Outgroup), genes used in the inference of each phylogeny (Genes), and the genomic compartment sampled by the genes used to infer each phylogeny (Genome). *Reimer et al., 2007b; Fig. 4.

Of the seven published zoanthidean trees that rely upon more than a single gene to support its inferences, only three (or 2.8% of the total) include attempted species trees based on analysis of concatenations (See section 3.1) of more than two genes originating from more than one genomic compartment (Table 1); even though this higher-level of genomic sampling (i.e., multiple genes from multiple genomic compartments) is the standard minimum practice in molecular phylogenetics and is rapidly being overshadowed by the use of hundreds of genes or whole genomes as the basis of phylogentic reconstructions (phylogenomics). These three analyses include the 6 gene, 93 species (representing 11 genera across 5 families) tree of Swain (2010), the 3 gene, 29 species (representing 12 genera across 3 families) tree of Sinniger et al. (2013), and the 5 gene, 48 species (representing 14 genera across 2 families) tree of Montenegro et al (2016) (Table 1). None of these analyses are as comprehensive as is currently possible (Table S1), both in terms of data available for taxa (at least 144 species, 25 genera, 9 families) and genes (at least 6 genes). Additionally, analyses that include genes with divergent hypervariable regions (e.g. mitochondrial 16S or nuclear ITS; see sections 3.2–3.3) either ignore these data (e.g., Montenegro et al., 2016) or limit the scope of research to closely related species (e.g., Sinniger et al., 2013) to eliminate difficult homology assessments (but, see Swain, 2009, 2010). With the spectacular proliferation of gene trees and discarded datasets (see sections 3.2–3.3), along with poor representation of species and higher taxa (less than 65% of species and 56% of genera and families for which there is available sequence for at least two genes are represented in any one of the previously published phylogenies), a comprehensive revision of the phylogeny of Zoanthidea seems warranted. Here, I present an updated comprehensive phylogeny for Zoanthidea and demonstrate that the use of a staggered sequence alignment to retain the evolutionary signal contained in hypervariable genes results in a more robust and defensible topology.

## 2. Materials and Methods

The phylogeny of Zoanthidea was revised to be current and comprehensive through maximum-likelihood inference applied to a concatenated multi-gene, multi-genomic alignment (see section 3.1). Nucleotide sequences of all unique Zoanthidea taxa, for which at least two different genes were available from GenBank, were included in the analysis (Table S1). Several of these genes contain hypervariable regions that are commonly used to differentiate species, but are also usually discarded for phylogenetic inference because of challenging homology assessments (see section 3.2). The effect of including hypervariable regions in tree inference (see section 3.3) was assessed through paired analyses: beginning with a reconstruction based on a staggered alignment that included every available nucleotide (see section 3.2, Fig. 1), followed by a reconstruction based on the identical alignment with an exclusion set to remove hypervariable positions and retain only conserved, universally alignable regions.

**Figure 1.**
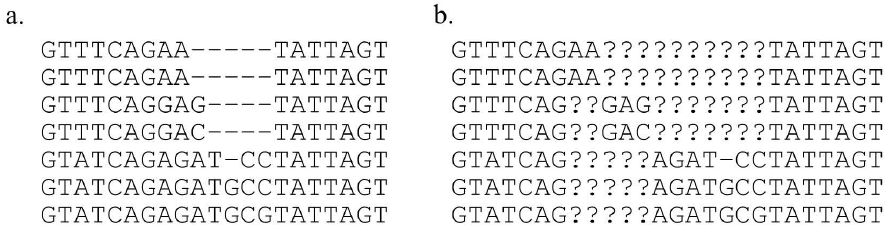
Example matrices aligned by standard (a) and staggered (b) protocols. Note that the standard alignment is “over-aligned”, assuming homology where it is unlikely.

### 2.1 Sampling strategy

All Zoanthidea taxa with at least two different nucleotide sequences available in GenBank for the six most commonly applied genes were included in the analyses. These genes include nuclear 18S, ITS, and 28S ribosomal RNA (rRNA) of the internal transcribed spacer (ITS) region and mitochondrial 12S and 16s rRNA and protein-coding COI (Table S1). Sequences were compiled by first targeting additional accessions available for the taxa used in the alignment of Swain (2010) (TreeBASE: S10492), and then expanding to include additional available taxa (that met the above criteria). Outgroup taxa were selected following Rodríguez et al. (2014) to include closely related *Relicanthus daphneae* and Antipatharia species and more distantly related Actiniaria species.

Identification of archived nucleotide sequences originating from unique species of Zoanthidea is challenging because of the rapid evolution of current taxonomic designations, the common practice of including unnamed and undescribed specimens without unique identifiers in previous phylogenetic analyses, use of molecular parataxonomy uncoupled from accepted species concepts for naming new taxa (see Mayden 1997 for a review of species concepts), and misidentification of taxa or substitution of contaminate sequences (in place of targeted taxa) within nucleotide databases (see Swain, 2010; Swain et al., 2015; Swain et al., 2016; Swain and Swain, 2014; and Table S1 for details of name usage tracking and examples of mislabeled accession identification). As a result, a simple search of GenBank for Zoanthidea will return a hodgepodge collection of intractable or incorrect identification labels attached to > 2,800 accessions (last accessed on May 30, 2016; http://www.ncbi.nlm.nih.gov/nuccore).

The only way to correctly identify which nucleotide sequences originate from the same specimens, and identify all available unique species (that meet the minimum genomic sampling detailed above), is to follow the usage of individual sequences through the 46 publications (and their associated supplemental files) and 107 molecular phylogenies that have been constructed for Zoanthidea (Table 1, Table S1). This required tracking GenBank accessions, publication-specific taxon and specimen names, and in some cases creating species specific nucleotide alignments to compare intraspecific specimens and identify mislabeled sequences (e.g. Swain and Swain, 2014).

### 2.2 Sequence alignment

A concatenated alignment was assembled by integrating additional sequences into the existing staggered framework of a previously published multiple alignment for Zoanthidea. This included every nucleotide of all the sequences analyzed (complete-data) and is the staggered alignment (See Table S1 for the genes and gene segments used for each species). Gene sequences originating from GenBank were edited to remove amplification primers and single nucleotide insertions from protein coding genes. Sequences were added to the alignment of Swain (2010) (TreeBase: S10492) and aligned manually in Bioedit 7.2.5 (Hall, 1999) following the existing staggered framework for hypervariable regions ITS1 and ITS2, and hypervariable regions within 16S and 12S (see section 3.2 for a description of theory, usage, and structure of staggered alignments and Table 2 for the alignment positions that were staggered). The sequences of the remaining three genes (18S, 28S, & COI), one gene segment (5.8S), and conserved regions of 16S and 12S conform to the format of a standard universal alignment within the complete-data staggered alignment. The standard alignment is derived directly from the staggered alignment by excluding the staggered hypervariable regions (see section 2.3 and Table 2).

**Table 2.**
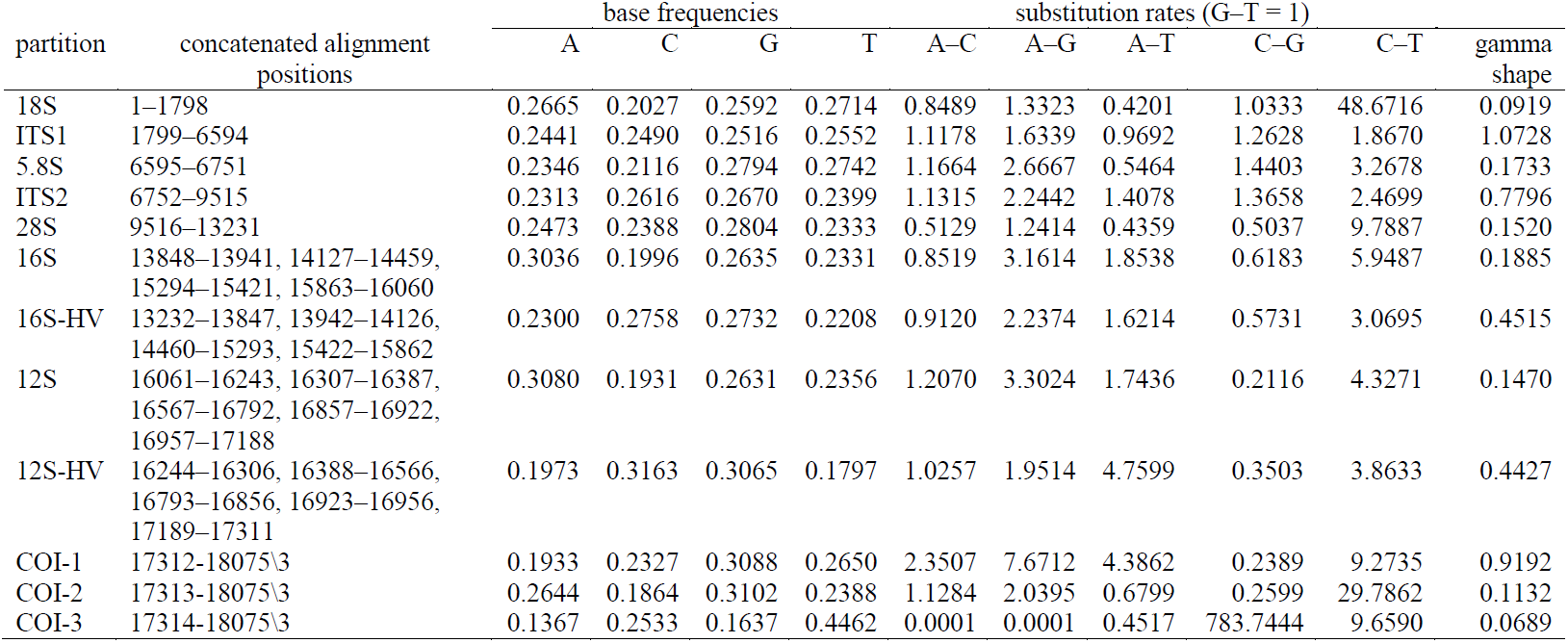
Partition definitions and per-partition parameter estimates used to model sequence evolution for phylogenetic inference. Hypervariable regions that were staggered in the staggered alignment, or excluded in the standard alignment, are alignment partitions ITS1, ITS2, 16S-HV, and 12S-HV.

| partition | concatenated alignment positions | base frequencies |  |  |  | substitution rates (G-T = 1) |  |  |  |  | gamma shape |
| --- | --- | --- | --- | --- | --- | --- | --- | --- | --- | --- | --- |
|  |  | A | C | G | T | A-C | A-G | A-T | C-G | C-T |  |
| 18S | 1-1798 | 0.2665 | 0.2027 | 0.2592 | 0.2714 | 0.8489 | 1.3323 | 0.4201 | 1.0333 | 48.6716 | 0.0919 |
| ITS1 | 1799-6594 | 0.2441 | 0.2490 | 0.2516 | 0.2552 | 1.1178 | 1.6339 | 0.9692 | 1.2628 | 1.8670 | 1.0728 |
| 5.8S | 6595-6751 | 0.2346 | 0.2116 | 0.2794 | 0.2742 | 1.1664 | 2.6667 | 0.5464 | 1.4403 | 3.2678 | 0.1733 |
| ITS2 | 6752-9515 | 0.2313 | 0.2616 | 0.2670 | 0.2399 | 1.1315 | 2.2442 | 1.4078 | 1.3658 | 2.4699 | 0.7796 |
| 28S | 9516-13231 | 0.2473 | 0.2388 | 0.2804 | 0.2333 | 0.5129 | 1.2414 | 0.4359 | 0.5037 | 9.7887 | 0.1520 |
| 16S | 13848-13941, 14127-14459, 15294-15421, 15863-16060 | 0.3036 | 0.1996 | 0.2635 | 0.2331 | 0.8519 | 3.1614 | 1.8538 | 0.6183 | 5.9487 | 0.1885 |
| 16S-HV | 13232-13847, 13942-14126, 14460-15293, 15422-15862 | 0.2300 | 0.2758 | 0.2732 | 0.2208 | 0.9120 | 2.2374 | 1.6214 | 0.5731 | 3.0695 | 0.4515 |
| 12S | 16061-16243, 16307-16387, 16567-16792, 16857-16922, 16957-17188 | 0.3080 | 0.1931 | 0.2631 | 0.2356 | 1.2070 | 3.3024 | 1.7436 | 0.2116 | 4.3271 | 0.1470 |
| 12S-HV | 16244-16306, 16388-16566, 16793-16856, 16923-16956, 17189-17311 | 0.1973 | 0.3163 | 0.3065 | 0.1797 | 1.0257 | 1.9514 | 4.7599 | 0.3503 | 3.8633 | 0.4427 |
| COI-1 | 17312-18075\3 | 0.1933 | 0.2327 | 0.3088 | 0.2650 | 2.3507 | 7.6712 | 4.3862 | 0.2389 | 9.2735 | 0.9192 |
| COI-2 | 17313-18075\3 | 0.2644 | 0.1864 | 0.3102 | 0.2388 | 1.1284 | 2.0395 | 0.6799 | 0.2599 | 29.7862 | 0.1132 |
| COI-3 | 17314-18075\3 | 0.1367 | 0.2533 | 0.1637 | 0.4462 | 0.0001 | 0.0001 | 0.4517 | 783.7444 | 9.6590 | 0.0689 |

Hypervariable regions are divergent in both sequence identity and length and are only alignable with homologous sequences from closely-related species. The staggered regions of the matrix are a form of phylogeny-informed alignment, where sequences are added to the matrix proximal to closely-related taxa that were previously aligned (Morrison et al., 2015). This allows visual detection and alignment of short sequences within hypervariable regions that are homologous in closely-related species, which are then isolated from divergent taxa within the matrix by inserting coding for unknown character states (question marks) in parallel matrix positions (see section 3.2). This creates a block of unambiguously aligned sequences and moves all divergent sequences further along the alignment, where the process is then repeated, resulting in the staggering of hypervariable regions. The initial assessment of sequence similarity is performed at the 5’ terminus of ITS1, and is therefore dependent on the evolution of sequences in this region; however similarity is continuously reassessed while proceeding in the 3’ direction and individual sequences may be aligned with different groups of species along the length of the alignment (all of which is completely documented in the TreeBase alignment accession: S20129). Using the previous framework (’jump-starting’: Morrison, 2006) greatly simplifies the process as homologous sequences are already identified and adding new taxa requires relatively few adjustments.

### 2.3 Model parameters and phylogenetic analyses

The complete-data staggered alignment was partitioned for model-fitting and phylogenetic analysis, and species trees were independently inferred from the staggered (complete-data) and standard (hypervariable-excluded) alignments. Partitioning traced the boundaries of ribosomal subunits, staggered hypervariable regions, and codon positions (following Li et al., 2008) for model-fitting and phylogenetic analyses (Table 2), however branch length optimization was linked during tree inference due to incomplete per-partition taxon sampling. Parameters of nucleotide evolution of the General Time Reversible (GTR) model with gamma (+C) were determined simultaneously to phylogenetic inference using maximum-likelihood analysis of RAxML v8.2.8 (Stamatakis, 2014) in the CIPRES Science Gateway v3.3 (Miller et al., 2010) for the complete-data staggered alignment, and for the hypervariable exclusion set applied to the same alignment (i.e., the standard alignment). This exclusion set followed partitions applied to hypervariable regions (Table 2) and was used to assess the effect of the staggered, rapidly evolving data (see section 3.3). Nonparametric bootstrap support was estimated in RAxML using GTR and a categorical per-site rate heterogeneity approximation (CAT) from 1000 pseudoreplicates (Stamatakis, 2014). Edwardsiid Actiniaria species were designated as the outgroup.

## 3. Theory

### 3.1 Gene trees and species trees

We have known, for more than 30 years, the critical distinction between gene trees and species trees (Pamilo and Nei, 1988). Gene trees, or phylogenetic inferences based upon a single gene, reconstruct the relationships among homologous variants of a gene sampled from different species and describes gene evolution. Species trees, or phylogenetic inferences based upon multiple genes or data sources, reconstruct the evolutionary histories of species and describes species evolution. Although both may use DNA sequence evolution as the basis of inference, gene evolution is not equivalent to species evolution (Pamilo and Nei, 1988), and analyses of species-level questions (e.g. species detection, identification, and systematics) based upon gene trees falsely assume equivalence in the evolution of genes and species.

This disparity between gene and species trees has multiple biological causes, but is ultimately due to the incomplete history of species that is provided by sampling a small proportion of their genomes. Evolutionary events such as gene deletion or duplication and horizontal gene transfer can cause dramatic differences in the evolutionary histories of genes and species, but usually occur under specific circumstances, regions of the genome, or evolutionary lineages (Edwards, 2009). Conversely, differing evolutionary rates among genes, resulting in incomplete lineage sorting (ILS), or deep coalescence, and the related issues of branch length heterogeneity and heterotachy, appear to be universal issues that affect all genes, genomes (organelle or nucleus), and lineages (Edwards, 2009).

There are various approaches to reconstructing species trees, but all attempt to do so by incorporating multiple regions of the genome, and if possible multiple genomes, into the same tree inference. The two main approaches involve concatenation of multiple genes into the same alignment which is then used to infer a species tree, and alternatively inferring a tree from each gene which are then used to infer a species tree under a coalescence model. Which is the best approach for obtaining a highly supported and statically consistent inference is currently a source of active debate, as each approach makes specific assumptions and carry specific weaknesses (Edwards, 2009; Roch and Warnow, 2015; Springer and Gatesy 2016). There is a potential for concatenated analysis to return statistically inconsistent trees because of discordance in gene evolution (Degnan and Rosenberg, 2006; Rosenberg, 2013), which species tree-estimation based on gene trees can overcome (even in the anomaly zone; Liu et al., 2010); however discordance between Zoanthidea phylogenies based on nuclear and mitochondrial genes has been previously demonstrated to be insignificant (Swain, 2010). This assessment was performed at the genome-level rather than the gene-level because of incomplete taxon sampling of each gene, making comparable gene trees possible for only a few taxa. Additionally, datasets with incomplete taxon sampling (such as the case here) are predicted to be inferred with greater confidence using concatenation methods (Edwards, 2009). Regardless of the approach, incorporation of a greater volume and diversity of data into the inference should improve resolution and confidence in comparison to gene trees.

### 3.2 *Staggered Alignments*

Multiple sequence alignment and tree inference are both critical to molecular phylogenetic analysis, however the quality and data content of tree inference is entirely dependent upon the quality and data content of alignment, as a phylogeny is a representation and interpretation of its underlying homology statements (Morrison, 2009a). Given the critical role of alignment as the primary homology assessment in phylogenetic inference, standard automated nucleotide sequence alignment is adept at assessing substitution events, but generally inadequate for generating homology hypotheses for most other common evolutionary events (e.g., deletion, duplication, insertion, inversion, and translocation; Morrison, 2009a). This limitation is generally not problematic for protein-coding regions because much of the evolutionary change observed in these sequences involves single residue substitutions and standard automated alignment based on sequence similarity can adequately assign homology within these events. However, only 1–20% of the genome of multicellular eukaryotes are composed of these conserved coding regions (Szymanski et al., 2007), leaving most genetic material and its evolutionary events not assessable by standard automated alignment (Morrison, 2009b). Additionally, many genes commonly applied to phylogenetic questions are not protein-coding and therefore most standard automated alignments of these genes contain misaligned sequences.

The main concern is over-alignment, or aligning sequences that are unlikely to be homologous. This issue is usually addressed in a standard alignment by excluding sequence regions with alignment challenges that are easily observable; meaning that even automated alignment requires manual correction to exclude low quality homology assessments (Morrison, 2009b). There are multiple alternative approaches to refine alignments depending upon the molecular function of the nucleotides involved (such as rRNA secondary structure prediction coupled with alignment correction or molecular morphometric analysis: e.g., Aguilar and Reimer, 2010; Swain and Taylor, 2003; Torres-Suarez, 2014), but all are focused on the same goal: retaining within the analysis as much information as possible (Morrison, 2009b).

One approach, which was commonly used prior to the wide-spread application of automated sequence alignment (Morrison, 2006), is the staggered alignment (Barta, 1997). A staggered alignment is a mosaic of local and universal nucleotide sub-alignments that are partially informed by phylogeny which allow the retention of all nucleotides (complete-data) without aligning non-homologues sequences (Morrison et al., 2015). This approach creates a single matrix that aligns homologous nucleotides within each sub-alignment: universal sub-alignments (equivalent to a standard alignment) are composed of conserved regions of the genome and include all taxa in the analysis, while local sub-alignments are composed of hypervariable regions of the genome and include only closely related taxa in each sub-alignment which are then staggered relative to other local sub-alignments (Fig. 1). A standard alignment can be derived directly from a staggered alignment by excluding the staggered local sub-alignments.

Like most standard automated alignment approaches, the staggered alignment relies upon sequence similarity to assess alignment quality, however it also benefits from the ability to examine multiple adjacent positions to specifically accommodate length variation among sequences. This approach can be initiated through automated alignment as a first approximation, but is completed through manual adjustment of nucleotide positioning following a single rule: do not align sequences that are not obviously homologous (Morrison, 2006). The resulting staggered alignment explicitly addresses our inability to simultaneously assess homology of complex evolutionary events across all targeted taxa, without discarding informative data. A standard alignment would follow the identical procedure of automated alignment as a first approximation which is then completed through manual adjustment; however with a standard alignment, the manual adjustment discards hypervariable sequence data rather than making it available for phylogenetic inference. Phylogenetic trees inferred from the same DNA sequences using staggered alignments will differ from trees inferred from standard alignments in resolution, branch order, and node support within the terminal clades, but the internal order of cladogenesis will be similar because they rely upon similar data matrices (Barta 1997).

### 3.3 *Use of hypervariable regions in phylogenetic reconstruction*

The need for molecular markers that are informative at species and population levels has driven interest in capturing information contained in hypervariable regions of genomes. There is broad consensus that these regions contain informative variation, and they are widely used as species-level markers in many systems (Coleman, 2007; Forsman et al., 2009). Although mitochondrial outpaces nuclear sequence evolution in most organisms (Creer, 2007), mitochondrial sequence evolution is extraordinarily slow among anthozoans and is often invariant among its most closely-related species (Shearer et al., 2002). The nuclear ribosomal ITS region contains a mosaic of secondary structural elements that can be relatively conserved (stems) or variable (loops and buldges) (Hillis and Dixon, 1991), and evolve under complex and varied evolutionary constraints. Hypervarible ITS sequences can differ in both length and nucleotide identity, which can cause challenges in assessing homology for multiple sequence alignment. Additionally, the ITS array can be repeated hundreds or thousands of times within a single genome and the exact sequence of nucleotides in each copy are homogenized to variying degrees of completion and precision by concerted evolution (Elder and Turner, 1995; Hillis and Dixon, 1991); allowing for potential intragenomic variation.

These two issues, alignment challenges and intragenomic variation, are widely acknowledged as being the primary obstacles to sucessfully using hypervariable ITS sequences in phylogentic inference. Alignment challenges can be mitigated by exclusively comparing regions with similar sequence length and identity through a staggered alignement (see section 3.2), thereby increasing the probablity of correct homology assesment at each matrix position. Intragenomic variation can be a more insidous problem, however intragenomic heterogeneity that is sufficient to mask evolutionary singnal is often confined to specific taxonomic groups (Coleman, 2003, 2007). For example, within the Anthozoa order Scleractinia, ITS sequences have been broadly applied to phylogenetics (reviewed in Forsman et al., 2009; Kitahara et al., 2016), and extreme intragenomic heterogeneity is largely confined to the genus *Acropora* (Wei et al., 2006). Although intragenomic variation is not a common target for analysis within Zoanthidea, direct sequencing of ITS regions results in largely unambigous sequence reads (suggesting that most copies are homogenized; e.g., Swain, 2009, 2010) and, if intragenomic heterogeneity is present, it appears to be generally insufficient to mask the evolutionary signal of species boundaries (but see Reimer et al., 2007c for example of hybridization) and intraspecific relationships (Aguilar and Reimer, 2010; Fujii and Reimer, 2011; Hibino et al., 2014; Irei et al., 2015; Kise and Reimer, 2016; Montenegro et al., 2016; Montenegro et al., 2015; Reimer et al., 2012a; Reimer and Fujii, 2010; Reimer et al., 2013a; Reimer et al., 2010c; Reimer et al., 2012b; Reimer et al., 2014; Reimer et al., 2008a; Reimer and Sinniger, 2010b; Reimer et al., 2008c; Reimer et al., 2007b; Risi and Macdonald, 2015; Risi and Macdonald, 2016; Sinniger et al., 2010a; Swain, 2009, 2010).

## 4 Results and Discussion

### 4.1 *Nucleotide alignment matrix*

The search of GenBank for novel taxon and gene sequences added 234 accessions (from 97 species) to the 273 accessions (from 82 species) retained from the alignment of Swain (2010), for a total of 767 genes or gene segments from 144 Zoanthidea and 11 outgroup species, including representatives of nearly all known Zoanthidea genera (sequences of Epizoanthidae genera *Paleozoanthus* and *Thoracactis* are unavailable; Table S1). This includes 114 accessions of COI; a very short sequence (<600 nt) that codes for a subunit of an enzyme of the electron transport chain. As in other Anthozoa taxa, COI has long been known to be nearly useless for phylogenetic inference among closely-related species as its evolutionary rate is >100 times slower than most marine invertebrates (Hellberg, 2006; Shearer et al., 2002; Stampar et al., 2012). These COI sequences are included here only because these data are available, not to encourage continued investment in their collection, which should be seen as a misuse of limited resources expended upon an uninformative marker. The remaining 7 gene segments targeted in this analysis are incompletely sampled, such that the matrix contains 653 Zoanthidea genes or gene segments out of a possible 1008 (144 taxa x 7 gene segments). While incomplete data matrices are not desirable, maximum likelihood phylogenetic analyses are generally robust to missing data issues and including incompletely sampled genes generally increases their accuracy relative to excluding them (Jiang et al., 2014; Streicher et al., 2016). Collection of the missing sequences and completion of this matrix, along with bolstering data for species that have available sequences but did not meet the minimal requirements for inclusion here (e.g.*Isozoanthus sulcatus*), should be a focus of future research on improving the resolution and confidence in the Zoanthidea phylogeny. Completing this matrix will also allow a more comprehensive analysis of potential gene tree discordance then is currently possible, as well as inferring statistically consistent species trees under a coalescence model if discordance is detected.

Once compiled, and properly staggered, the alignment contains >2.8 million matrix positions (155 rows by 18,075 columns; TreeBase: S20129). Alignment staggering represents a significant time commitment over automated multiple alignment alone, as hypervariable regions of small subsamples of taxa are individually assessed and manually adjusted. However, the resulting staggered alignment simultaneously maximizes information content and analysis power as it retains all nucleotides recovered from the original sequencing reads and allows the application of nucleotide evolutionary model-informed phylogenetic inference. There are alternative methods available, such as secondary structure prediction followed by alignment correction or molecular morphometric analysis (e.g., Aguilar and Reimer, 2010; Swain and Taylor, 2003; Torres-Suarez, 2014), but all retain less of the original sequence data or restrict the diversity of taxa that can be simultaneously analyzed and employ less powerful and robust analyses. Partitioning the data matrix resulted in twelve distinct models of nucleotide evolution, detailed in Table 2, that were applied in ML inferences of tree topology and bootstrap support.

### 4.2 *Complete-data phylogenetic inference: analysis of the staggered alignment*

A search for the optimal ML tree from the partitioned staggered alignment resulted in a best tree with a likelihood score of −65296 (Fig. 2). This analysis recovered highly supported (bootstrap values of 100) monophylies of taxa representing order Zoanthidea and suborder Brachycnemina, with suborder Macrocnemina ancestral to Brachycnemina, and an overall tree topology that generally conforms to previously reported phylogenies that are of comparable taxon sampling (i.e. Sinniger et al., 2005; Swain, 2010).

**Figure 2.**
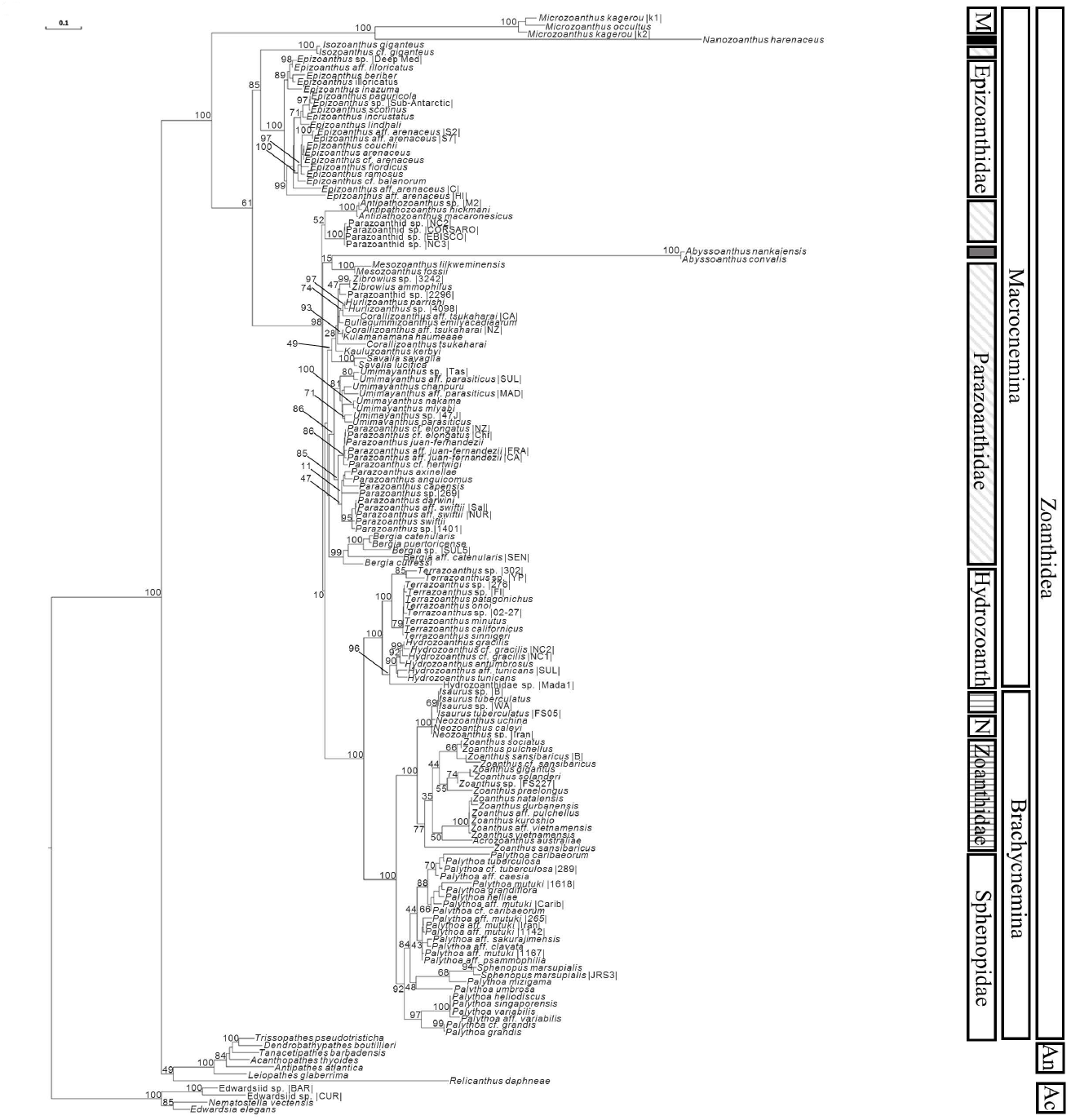
Complete-data tree. Maximum likelihood phylogeny of Zoanthidea based on a staggered alignment of concatenated nuclear (18S, ITS1, 5.8S, ITS2, & 28S) and mitochondrial (12S & 16S) ribosomal RNA and mitochondrial protein-coding (COI) nucleotide sequences. Support indicated by 1000 pseudoreplicate maximum likelihood bootstrap values. Taxonomic notations: order, Ac = Actiniaria, An = Antipatharia; family, M = Microzoanthidae, black bar = Nanozoanthidae, gray diagonal lines = Parazoanthidae, gray bar = Abyssoanthidae, Hydrozoanth = Hydrozoanthidae, gray horizontal line = Zoanthidae, N = Neozoanthidae.

Most currently recognized families were also recovered at a high-level of certainty including Microzoanthidae, Epizoanthidae, Abyssoanthidae, Hydrozoanthidae, and Sphenopidae. Nanozoanthidae is monospecific and therefore impossible to assess. Parazoanthidae is both paraphyletic (with respect to Abyssoanthidae) and polyphyletic (with respect to *Isozoanthus*) and the relationships among genera are largely unresolved (i.e., extremely weak bootstrap support) except for monophyletic sister genera *Parazoanthus* and *Umimayanthus*. Other monophyletic Parazoanthidae genera include *Antipathozoanthus, Mesozoanthus, Zibrowius, Hurlizoanthus, Savalia*, and *Bergia*. *Corallizoanthus* is polyphyletic, and *Bullagummizoanthus, Kauluzoanthus,* and *Kulamanamana* are monospecific (and therefore impossible to assess); however the topology in this region of the tree generally lacks support due to the paucity of sequence data available for these taxa (which is almost entirely mitochondrial; Table S1). Clarification of these relationships should be a priority for future research and could be easily accomplished by adding highly informative ITS and 28S genes. Zoanthidae is paraphyletic with respect to Neozoanthidae, where the genus *Neozoanthus* is ancestral to *Isaurus*. Also within Zoanthidae, *Zoanthus* is paraphyletic with respect to *Acrozoanthus*. Although Sphenopidae is monophyletic, its daughter genus *Palythoa* is paraphyletic with respect to *Sphenopus*. Although there is much more confidence in this region of the tree, ˜37% of the brachycnemic taxa are represented exclusively by mitochondrial genes in the data matrix and further resolution and confidence could be obtained by completing the matrix with information derived from the nuclear compartment.

### 4.3 *Hypervariable-excluded phylogenetic inference: analysis of the standard alignment*

A search for the optimal ML tree from the partitioned standard alignment, (equivalent to the staggered alignment excluding hypervariable regions; Table 2), resulted in a best tree with a likelihood score of −31899 (Fig. 3). Although the exact branching order of the two trees varies, the overall toplology of the hypervariable-excluded tree is largely congruent with the complete-data tree, and apart from a few exceptions matches the monophylies of higher taxa detailed above. The hypervariable-excluded tree differs with a polyphyletic Sphenopidae, *Abyssoanthus* within *Parazoanthus, Acrozoanthus* basal to *Zoanthus* and *Zoanthus* as monophyletic, and *Sphenopus* basal to Brachycnemina. Additionally, the bootstrap values are ˜15% lower in the hypervariable-excluded tree, with comparable nodes falling from a mean of 80.1 in the complete-data tree to 68.5 in the hypervariable-excluded tree.

**Figure 3.**
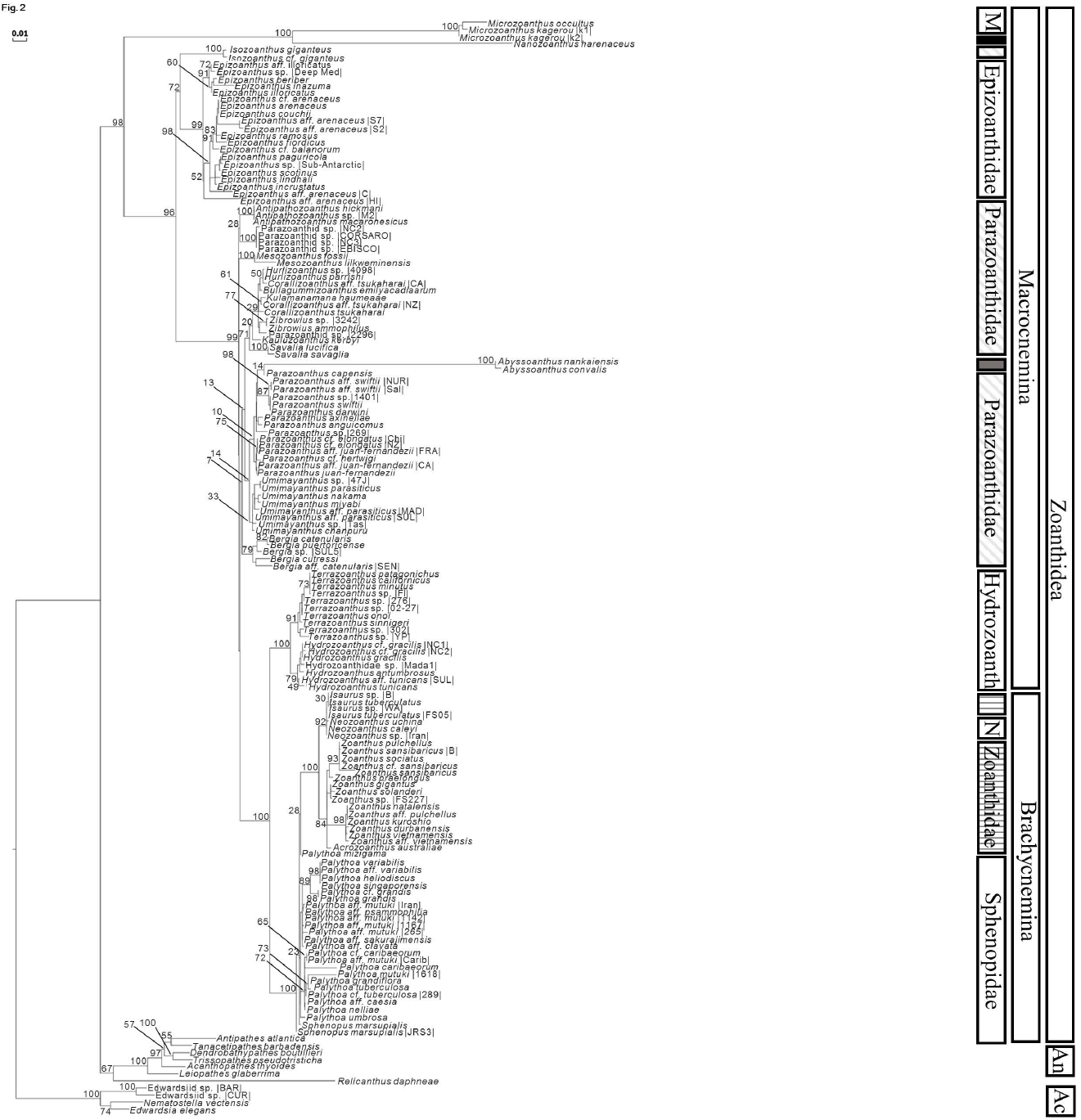
Hypervariable-excluded tree. Maximum likelihood phylogeny of Zoanthidea based on standard alignment of concatenated nuclear (18S, 5.8S, & 28S) and mitochondrial (12S & 16S) ribosomal RNA and mitochondrial protein-coding (COI) nucleotide sequences, with hypervariable regions of 12S & 16S (12S-HV, 16S-HV; Table 2) excluded. Support indicated by 1000 pseudoreplicate maximum likelihood bootstrap values. Taxonomic notations: order, Ac = Actiniaria, An = Antipatharia; family, M = Microzoanthidae, black bar = Nanozoanthidae, gray diagonal lines = Parazoanthidae, gray bar = Abyssoanthidae, Hydrozoanth = Hydrozoanthidae, gray horizontal line = Zoanthidae, N = Neozoanthidae.

### 4.4 *Preferred topology: the complete-data inference based on the staggered alignment*

Given these competing topologies, the complete-data tree based on the staggered alignment best reflects our current understanding of molecular evolution, evolution of form, and systematics of Zoanthidea. The position of *Abyssoanthus* is generally problematic (and weakly supported in both inferences) because its nucleotide sequences are highly divergent (and therefore rarely included in phylogenetic analyses) and its anatomy is all but unknown (Reimer and Sinniger, 2010a; Reimer et al., 2007a); however, *Abyssoanthus* as sister to *Mesozoanthus* (as in the complete-data tree based on the staggered alignment) agrees with previous phylogenetic hypotheses (Reimer and Sinniger, 2010a; Reimer et al., 2007a) and the ecology of both taxa, whereas *Abyssoanthus* within *Parazoanthus* (as in the hypervariable-excluded tree based upon the standard alignment) would be a significant departure from our understanding of both genera.

In the hypervariable-excluded tree (based upon the standard alignment), Sphenopidae is polyphyletic because *Palythoa mizigama* is inferred to be part of the Zoanthidae monophyly. Zoanthidae and Sphenopidae have contrasting anatomical features (reviewed in Swain et al., 2016) and there is no indication that the morphology of *P. mizigama* differs from our understanding of *Palythoa* and Sphenopidae (Irei et al., 2015). *Acrozoanthus* and *Sphenopus* are both basal to their respective clades in the hypervariable-excluded tree (based upon the standard alignment), supporting the validity of these genera; however, both are odd genera represented by few species that might best fit within other genera, as inferred in the complete-data tree (based upon the staggered alignment). *Acrozoanthus* is a mono-specific genus that was created because of its apparent ability to build an erect skeleton (Saville-Kent, 1893), however this skeleton was revealed to be the parchment-like tube of a worm in the genus *Eunice* (Ryland, 1997). Therefore skeleton-building is not the odd character of *Acrozoanthus*, rather is its ability to form symbiotic associations with polycheate worms (which is only common among zoanthideans in the genus *Epizoanthus* (Swain, 2010, Reimer et al., 2010a) and cannot help us to understand the relationship between *Acrozoanthus* and *Zoanthus*). Including *Acrozoanthus* within *Zoanthus* has been previously suggested using morphology (reviewed in Ryland, 1997) and molecular phylogenetics (e.g. Reimer et al., 2010c; Sinniger et al., 2005; Swain, 2010), and the complete-data tree (based upon the staggered alignment) agrees with this conclusion. *Sphenopus* is odd because it is solitary and azooxanthellate, whereas most of Sphenopidae are colonial and zooxanthellate. However, two recently described *Palythoa* species, *P. umbrosa* and *P. mizigama*, form colonies with small numbers of polyps and are uncharacteristically azooxanthellate (Irei et al., 2015). *Sphenopus* is part of a monophyly with *P. umbrosa* and *P. mizigama* within the *Palythoa* monophyly in the complete-data tree (based upon the staggered alignment), but these taxa are dispersed across the hypervariable-excluded tree (based upon the standard alignment). All of these major differences between these two trees suggest that the topology of the complete-data tree (based upon the staggered alignment) is a more accurate representation.

### 4.5 *Novel hypotheses*

Although the complete-data tree (based upon the staggered alignment) is largely congruent with previous comparable phylogenies, it also supports novel hypotheses and discredits others. Previous work on Microzoanthidae and Nanozoanthidae had placed these families within a monophyly with the Parazoanthidae genus *Isozoanthus* and as sister to the remaining Zoanthidea (Fujii and Reimer, 2011, 2013). The complete-data tree (based upon the staggered alignment) and the hypervariable-excluded tree (based upon the standard alignment) strongly supports the hypothesis that Microzoanthidae and Nanozoanthidae form an exclusive monophyly at the base of Zoanthidea and that *Isozoanthus* is sister to *Epizoanthus* (Fig. 2, 3).

The region of the complete-data tree (based upon the staggered alignment) containing the octocoral-symbiotic genera and their allies (*Bullagummizoanthus, Corallizoanthus, Hurlizoanthus, Kauluzoanthus, Kulamanamana, Savalia, Zibrowius*) lacks strong support, mostly because there are data for only 3 of 6 genes targeted in these analyses (Table S1), but suggests that many of these taxa could be congeneric. Unfortunately, along with the paucity of genetic data for these taxa, there is also very little anatomical data upon which a hypothesis could be based (Swain et al., 2016). There is considerable information about the genetics and anatomy of the three *Corallizoanthus* species included here (Swain, 2010; Swain et al., 2015; Swain et al., 2016; Swain and Swain, 2014), which could serve as guide for reexamination of the remaining taxa in this group. Resolving this portion of the tree should be a priority for future research.

At the base of the *Palythoa* monophyly is a strongly supported clade of taxa that is almost entirely comprised of species that were once assigned to the genus *Protopalythoa*. This includes *P. grandis, P. heliodiscus, P. variabilis*, but not others such as *P. grandiflora* and *P. mutuki*. The status of this genus has been in dispute for some time (Burnett et al., 1997, Low et al.,2016; Reimer et al., 2006c; Ryland and Lancaster, 2003), and the findings presented here do not settle this issue, but perhaps this novel hypothesis is an opening to reconsider the validity of *Protopalythoa*. Again, additional DNA sequence that could be used to complete the data matrix could dramatically improve our understanding of the taxa in this region.

### 4.6 *Sister of Zoanthidea*

Where the order Zoanthidea inserts into the Cnidaria Tree of Life has been one of the targets of sustained research and a point of contention (see Rodríguez et al., 2014 and references therein). Sister to Zoanthidea has been variously hypothesized to be either Actiniaria or Antipatharia, and its relationship with the remaining orders is poorly understood. The recent Actiniaria-focused inference by Rodríguez et al. (2014) put forth the novel hypothesis that the enigmatic species *Relicanthus daphneae* may be sister to Zoanthidea and both form a monophyly with Antipatharia, while Actinaria is much more distantly related. The Zoanthidea-focused analysis presented here included representatives of Antipatharia, Actiniaria, and *R. daphneae*, however the root of the phylogenetic inference was set only as the two undescribed Edwardsiidae species, allowing the remaining taxa to be placed according to the data. In this analysis *R. daphneae* is sister to Antipatharia and both form a monophyly with Zoanthidea (Fig. 1, 2). With the highly divergent sequences of *R. daphneae* and weak bootstrap support at its insertion, along with partially-overlapping taxon and datasets between the analysis presented here and Rodríguez et al. (2014), it would seem to remain an open question.

## 5 Conclusions

Proliferation of Zoanthidea species and higher taxa descriptions since the year 2000, have occurred at a rate not seen for nearly 100 years (Swain et al., 2016). This explosion of discovery and identification of evolutionary patterns was triggered by the application of molecular phylogenetics to an order that was largely ignored because rampant confusion at the species and genus levels. Molecular tools have detected cryptic species and genera, both reinforced and dismantled previous taxonomic and systematic hypotheses, and greatly expanded our understanding of Zoanthidea diversity. However, the tendency to, almost exclusively, construct gene trees based on many specimens of the same species (Table 1) as a tool of species discovery, has caused a parallel proliferation in published trees without a corresponding increase in the understanding of species phylogeny. The comprehensive phylogeny presented here is intended to fill that gap. Its utility is not in simply describing the proximate relationships of individual species as supported by the nucleotide data, as the phylogenies presented here are generally in agreement with previously published Zoanthidea species trees and many gene trees (but see section 4.5 for novel hypotheses). Rather the major advancement here is the inclusion of nearly all available species and nucleotide sequence data (including data that are usually discarded) into a single robust phylogeny (increasing resolution and node certainty) that can then serve as an expandable tool for performing higher-level assessments of evolutionary hypotheses and phylogeny-corrected statistical analyses (see exemplar analyses in Swain et al., 2015; Swain et al., 2016).

The absence of a comprehensive phylogeny can be traced to the disparity between the simplicity of alignment and the data content of the genes currently being applied to this group. Conserved genes can be easily aligned across taxa representing all genera and families of Zoanthidea, but produces poorly resolved and poorly supported trees. Hypervariable genes can provide the evolutionary signal necessary for topology resolution and confidence building, but creating the homology statements (sequence alignments) necessary for phylogenetic inference is a challenge. By staggering the alignment, it is possible to overcome the challenge of hypervariable sequence evolution while retaining its information. The complete-data tree based upon the staggered alignment provides a better supported topology that best matches our understanding of evolution among Zoanthidea taxa. All major incongruencies between the hypervariable-excluded (based upon the standard alignment) and complete-data (based upon the staggered alignment) trees favor the hypothesis offered by the complete-data tree.

## Acknowledgments

This work was supported by the US National Science Foundation [CBET-0937987, CBET-1240416, and OCE-0550599]; Northwestern University, Evanston, IL; the Field Museum of Natural History, Chicago, IL; and Florida State University, Tallahassee, FL; and L. Marcelino, and V. Backman of NU, R. Bieler of FMNH, and J. Wulff and S. Steppan of FSU. My original use of staggered alignments in phylogenetic analyses grew out of conversations on analyzing hypervariable sequences with S. Steppan.

---

Table S1.

Zoanthidea specimens and GenBank accession numbers for each of the sequences used for phylogenetic inference. Accessions of sequences not included in the alignment of Swain (2010) (TreeBASE: S10492) are in bold.

## References

Aguilar, C., Reimer, J.D., 2010. Molecular phylogenetic hypotheses of *Zoanthus* species (Anthozoa:Hexacorallia) using RNA secondary structure of the internal transcribed spacer 2 (ITS2). Marine Biodiversity 40, 195–204.

Barta, J.R., 1997. Investigating phylogenetic relationships within the *Apicomplexa* using sequence data: The search for homology. Methods-a Companion to Methods in Enzymology 13, 81–88.

Burnett, W.J., Benzie, J.A.H., Beardmore, J.A., Ryland, J.S., 1997. Zoanthids (Anthozoa, Hexacorallia) from the Great Barrier Reef and Torres Strait, Australia: Systematics, evolution and a key to species. Coral Reefs 16, 55–68.

Coleman, A.W., 2003. ITS2 is a double-edged tool for eukaryote evolutionary comparisons. Trends in Genetics 19, 370–375.

Coleman, A.W., 2007. Pan-eukaryote ITS2 homologies revealed by RNA secondary structure. Nucleic Acids Research 35, 3322–3329.

Creer, S., 2007. Choosing and using introns in molecular phylogenetics. Evolutionary Bioinformatics 3, 99–108.

Degnan, J.H., Rosenberg, N.A., 2006. Discordance of species trees with their most likely gene trees. PLoS Genetics, pp. 762–768.

Edwards, S.V., 2009. Is a new and general theory of molecular systematics emerging? Evolution 63, 1–19.

Elder, J.F., Turner, B.J., 1995. Concerted evolution of repetitive DNA sequences in eukaryotes. Quarterly Review of Biology 70, 297–320.

Forsman, Z.H., Barshis, D.J., Hunter, C.L., Toonen, R.J., 2009. Shape-shifting corals: molecular markers show morphology is evolutionarily plastic in *Porites*. BMC Evolutionary Biology 9, 45.

Fujii, T., Reimer, J.D., 2011. Phylogeny of the highly divergent zoanthid family Microzoanthidae (Anthozoa, Hexacorallia) from the Pacific. Zoologica Scripta 40, 418–431.

Fujii, T., Reimer, J.D., 2013. A new family of diminutive zooxanthellate zoanthids (Hexacorallia: Zoantharia). Zoological Journal of the Linnean Society 169, 509–522.

Hall, T.A., 1999. BioEdit: a user-friendly biological sequence alignment editor and analysis program for Windows 95 /98 /NT. Nucleic Acid Symposium Series, pp. 95–98.

Hellberg, M.E., 2006. No variation and low synonymous substitution rates in coral mtDNA despite high nuclear variation. BMC Evolutionary Biology 6, 24

Hibino, Y., Todd, P.A., Yang, S.-y., Benayahu, Y., Reimer, J.D., 2014. Molecular and morphological evidence for conspecificity of two common Indo-Pacific species of *Palythoa* (Cnidaria: Anthozoa). Hydrobiologia 733, 31–43.

Hillis, D.M., Dixon, M.T., 1991. Ribosomal DNA: molecular evolution and phylogenetic inference. Quarterly Review of Biology 66, 411–453.

Irei, Y., Sinniger, F., Reimer, J.D., 2015. Descriptions of two azooxanthellate *Palythoa* species (Subclass Hexacorallia, Order Zoantharia) from the Ryukyu Archipelago, southern Japan. Zookeys 478, 1–26.

Jiang, W., Chen, S.Y., Wang, H., Li, D.Z., Wiens, J.J., 2014. Should genes with missing data be excluded from phylogenetic analyses? Molecular Phylogenetics and Evolution 80, 308–318.

Kise, H., Reimer, J.D., 2016a. Unexpected diversity and a new species of *Epizoanthus* (Anthozoa, Hexacorallia) attached to eunicid worm tubes from the Pacific Ocean. Zookeys 562, 49–71.

Kitahara, M.V., Fukami, H., Benzoni, F., Huang, D., 2016. The new systematics of Scleractinia: integrating molecular and morphological evidence. In: Goffredo, S., Dubinsky, Z. (Eds.), The Cnidaria, Past, Present and Future: The world of Medusa and her sisters. Springer International Publishing, pp. 41–59.

Koupaei, A.N., Mostafavi, P.G., Mehrabadi, J.F., Fatemi, S.M.R., 2014. Molecular diversity of coral reef-associated zoanthids off Qeshm Island, northern Persian Gulf. International Aquatic Research 6, 64.

Koupaei, A.N., Mostafavi, P.G., Mehrabadi, J.F., Fatemi, S.M.R., Dehghani, H., 2016. Diversity of shallow water zoantharians in Hengam and Larak Islands, in the Persian Gulf. Journal of the Marine Biological Association of the United Kingdom 96, 1145–1155.

Li, C., Lu, G., Orti, G., 2008. Optimal data partitioning and a test case for ray-finned fishes (Actinopterygii) based on ten nuclear loci. Systematic Biology 57, 519–539.

Liu, L.A., Yu, L.L., Edwards, S.V., 2010. A maximum pseudo-likelihood approach for estimating species trees under the coalescent model. BMC Evolutionary Biology 10, 302.

Miller, M.A., Pfeiffer, W., Schwartz, T., 2010. Creating the CIPRES Science Gateway for inference of large phylogenetic trees. Gateway Computing Environments Workshop (GCE), New Orleans, LA, pp. 1–8.

Montenegro, J., Low, M.E.Y., Reimer, J.D., 2016. The resurrection of the genus *Bergia*(Anthozoa, Zoantharia, Parazoanthidae). Systematics and Biodiversity 14, 63–73.

Montenegro, J., Sinniger, F., Reimer, J.D., 2015. Unexpected diversity and new species in the sponge-Parazoanthidae association in southern Japan. Molecular Phylogenetics and Evolution 89, 73–90.

Morrison, D.A., 2006. Multiple sequence alignment for phylogenetic purposes. Australian Systematic Botany 19, 479–539.

Morrison, D.A., 2009a. A framework for phylogenetic sequence alignment. Plant Systematics and Evolution 282, 127–149.

Morrison, D.A., 2009b. Why would phylogeneticists ignore computerized sequence alignment? Systematic Biology 58, 150–158.

Morrison, D.A., Morgan, M.J., Kelchner, S.A., 2015. Molecular homology and multiple-sequence alignment: an analysis of concepts and practice. Australian Systematic Botany 28, 46–62.

Pamilo, P., Nei, M., 1988. Relationships between gene trees and species trees. Molecular Biology and Evolution 5, 568–583.

Reimer, J.D., Foord, C., Yuka Irei, Y., 2012a. Species diversity of shallow water zoanthids (Cnidaria: Anthozoa: Hexacorallia) in Florida. Journal of Marine Biology 2012, 856079.

Reimer, J.D., Fujii, T., 2010. Four new species and one new genus of zoanthids (Cnidaria, Hexacorallia) from the Galapagos Islands. Zookeys, 1–36.

Reimer, J.D., Hirose, M., Irei, Y., Obuchi, M., Sinniger, F., 2011a. The sands of time: rediscovery of the genus Neozoanthus (Cnidaria: Hexacorallia) and evolutionary aspects of sand incrustation in brachycnemic zoanthids. Marine Biology 158, 983–993.

Reimer, J.D., Hirose, M., Nishisaka, T., Sinniger, F., Itani, G., 2010a. *Epizoanthus* spp. associations revealed using DNA markers: A case study from Kochi, Japan. Zoological Science 27, 729–734.

Reimer, J.D., Hirose, M., Wirtz, P., 2010b. Zoanthids of the Cape Verde Islands and their symbionts: previously unexamined diversity in the Northeastern Atlantic. Contributions to Zoology 79, 147–163.

Reimer, J.D., Hirose, M., Yanagi, K., Sinniger, F., 2011b. Marine invertebrate diversity in the oceanic Ogasawara Islands: a molecular examination of zoanthids (Anthozoa: Hexacorallia) and their *Symbiodinium* (Dinophyceae). Systematics and Biodiversity 9, 133–143.

Reimer, J.D., Irei, Y., Fujii, T., Yang, S.Y., 2013a. Molecular analyses of shallow-water zooxanthellate zoanthids (Cnidaria: Hexacorallia) from Taiwan and their *Symbiodinium* spp. Zoological Studies 52, 38.

Reimer, J.D., Irei, Y., Naruse, T., 2013b. A record of *Neozoanthus cf. uchina* Reimer, Irei & Fujii, 2012 from the Yaeyama Islands, southern Ryukyu Islands, Japan. Fauna Ryukyuana 1–7.

Reimer, J.D., Ishikawa, S.A., Hirose, M., 2010c. New records and molecular characterization of *Acrozoanthus* (Cnidaria: Anthozoa: Hexacorallia) and its endosymbionts (*Symbiodinium* spp.) from Taiwan. Marine Biodiversity 41, 313–323.

Reimer, J.D., Lin, M., Fujii, T., Lane, D.J.W., Hoeksema, B.W., 2012b. The phylogenetic position of the solitary zoanthid genus *Sphenopus* (Cnidaria: Hexacorallia). Contributions to Zoology 81, 43–54.

Reimer, J.D., Lorion, J., Irei, Y., Hoeksema, B., Wirtz, P., 2014. Ascension Island shallow-water Zoantharia (Hexacorallia: Cnidaria) and their zooxanthellae (*Symbiodinium*). Journal of the Marine Biological Association of the United Kingdom doi:10.1017/S0025315414000654.

Reimer, J.D., Nonaka, M., Sinniger, F., Iwase, F., 2008a. Morphological and molecular characterization of a new genus and new species of parazoanthid (Anthozoa: Hexacorallia: Zoantharia) associated with Japanese Red Coral. Coral Reefs 27, 935–949.

Reimer, J.D., Ono, S., Fujiwara, Y., Takishita, K., Tsukahara, J., 2004. Reconsidering *Zoanthus* spp. diversity: Molecular evidence of conspecifity within four previously presumed species. Zoological Science 21, 517–525.

Reimer, J.D., Ono, S., Iwama, A., Takishita, K., Tsukahara, J., Maruyama, T., 2006a. Morphological and molecular revision of Zoanthus (Anthozoa: Hexacorallia) from southwestern Japan, with descriptions of two new species. Zoological Science 23, 261–275.

Reimer, J.D., Ono, S., Iwama, A., Tsukahara, J., Maruyama, T., 2006b. High levels of morphological variation despite close genetic relatedness between *Zoanthus aff. vietnamensis* and *Zoanthus kuroshio* (Anthozoa: Hexacorallia). Zoological Science 23, 755–761.

Reimer, J.D., Ono, S., Takishita, K., Tsukahara, J., Maruyama, T., 2006c. Molecular evidence suggesting species in the zoanthid genera *Palythoa* and *Protopalythoa* (Anthozoa: Hexacorallia) are congeneric. Zoological Science 23, 87–94.

Reimer, J.D., Ono, S., Tsukahara, J., Iwase, F., 2008b. Molecular characterization of the zoanthid genus *Isaurus* (Anthozoa: Hexacorallia) and associated zooxanthellae (*Symbiodinium* spp.) from Japan. Marine Biology 153, 351–363.

Reimer, J.D., Sinniger, F., 2010a. Discovery and description of a new species of *Abyssoanthus* (Zoantharia: Hexacorallia) at the Japan Trench: the world’s deepest known zoanthid. Cahiers De Biologie Marine 51, 451–457.

Reimer, J.D., Sinniger, F., 2010b. Unexpected diversity in Canadian Pacific zoanthids (Cnidaria: Anthozoa: Hexacorallia): a molecular examination and description of a new species from the waters of British Columbia. Marine Biodiversity 40, 249–260.

Reimer, J.D., Sinniger, F., Fujiwara, Y., Hirano, S., Maruyama, T., 2007a. Morphological and molecular characterisation of *Abyssoanthus nankaiensis*, a new family, new genus and new species of deep-sea zoanthid (Anthozoa: Hexacorallia: Zoantharia) from a north-west Pacific methane cold seep. Invertebrate Systematics 21, 255–262.

Reimer, J.D., Sinniger, F., Hickman, C.P., 2008c. Zoanthid diversity (Anthozoa: Hexacorallia) in the Galapagos Islands: a molecular examination. Coral Reefs 27, 641–654.

Reimer, J.D., Takishita, K., Ono, S., Maruyama, T., 2007b. Diversity and evolution in the zoanthid genus *Palythoa* (Cnidaria: Hexacorallia) based on nuclear ITS-rDNA. Coral Reefs 26, 399–410.

Reimer, J.D., Takishita, K., Ono, S., Tsukahara, J., Maruyama, T., 2007c. Molecular evidence suggesting interspecific hybridization in Zoanthus spp. (Anthozoa: Hexacorallia). Zoological Science 24, 346–359.

Reimer, J.D., Todd, P.A., 2009. Preliminary molecular examination of zooxanthellate zoanthids (Hexacorallia: Zoantharia) and associated zooxanthellae (*Symbiodinium* spp.) diversity in Singapore. Raffles Bulletin of Zoology 22, 103–120.

Risi, M.M., Macdonald, A.H.H., 2015. Possible synonymies of *Zoanthus* (Anthozoa: Hexacorallia) species on the east coast of South Africa with Pacific congeners. Systematics and Biodiversity 13, 93–103.

Risi, M.M., Macdonald, A.H.H., 2016. Molecular examination of rocky shore brachycnemic zoantharians (Anthozoa: Hexacorallia) and their *Symbiodinium* symbionts (Dinophyceae) in the southwest Indian Ocean. Marine Biodiversity 46, 113–127.

Roch, S., Warnow, T., 2015. On the robustness to gene tree estimation error (or lack thereof) of coalescent-based species tree methods. Systematic Biology 64, 663–676.

Rodríguez, E., Barbeitos, M.S., Brugler, M.R., Crowley, L.M., Grajales, A., Gusmão, L., Häussermann, V., Reft, A., Daly, M., 2014. Hidden among sea anemones: The first comprehensive phylogenetic reconstruction of the order actiniaria (Cnidaria, Anthozoa, Hexacorallia) reveals a novel group of hexacorals. PLoS One 9, e96998.

Rosenberg, N.A., 2013. Discordance of species trees with their most likely gene trees: a unifying principle. Molecular Biology and Evolution 30, 2709–2713.

Ryland, J.S., 1997. Budding in *Acrozoanthus* Saville-Kent, 1893 (Anthozoa: Zoanthidea). In: den Hartog, J. (Ed.), 6th International Conference on Coelenterate Biology. Nationaal Natuurhistorisch Museum, pp. 423–428.

Ryland, J.S., Lancaster, J.E., 2003. Revision of methods for separating species of *Protopalythoa* (Hexacorallia: Zoanthidea) in the tropical West Pacific. Invertebrate Systematics 17, 407–428.

Santos, M.E.A., Kitahara, M.V., Lindner, A., Reimer, J.D., 2015. Overview of the order Zoantharia (Cnidaria: Anthozoa) in Brazil. Marine Biodiversity 46, 547–559.

Saville-Kent, W., 1893. The Great Barrier Reef of Australia: its products and potentialities. Allen, W.H., London. pp. 1–387.

Shearer, T.L., Van Oppen, M.J.H., Romano, S.L., Worheide, G., 2002. Slow mitochondrial DNA sequence evolution in the Anthozoa (Cnidaria). Molecular Ecology 11, 2475–2487.

Shiroma, E., Reimer, J.D., 2010. Investigations into the reproductive patterns, ecology, and morphology in the zoanthid genus *Palythoa* (Cnidaria: Anthozoa: Hexacorallia) in Okinawa, Japan. Zoological Studies 49, 182–194.

Sinniger, F., Haussermann, V., 2009. Zoanthids (Cnidaria: Hexacorallia: Zoantharia) from shallow waters of the southern Chilean fjord region, with descriptions of a new genus and two new species. Organisms Diversity & Evolution 9, 23–36.

Sinniger, F., Montoya-Burgos, J.I., Chevaldonne, P., Pawlowski, J., 2005. Phylogeny of the order Zoantharia (Anthozoa, Hexacorallia) based on the mitochondrial ribosomal genes. Marine Biology 147, 1121–1128.

Sinniger, F., Ocana, O.V., Baco, A.R., 2013. Diversity of zoanthids (Anthozoa: Hexacorallia) on Hawaiian seamounts: Description of the Hawaiian gold coral and additional zoanthids. PLoS One 8, e52607.

Sinniger, F., Reimer, J.D., Pawlowski, J., 2008. Potential of DNA sequences to identify zoanthids (Cnidaria: Zoantharia). Zoological Science 25, 1253–1260.

Sinniger, F., Reimer, J.D., Pawlowski, J., 2010a. The Parazoanthidae (Hexacorallia: Zoantharia) DNA taxonomy: description of two new genera. Marine Biodiversity, 57–70.

Sinniger, F., Zelnio, K.A., Taviani, M., Reimer, J.D., 2010b. Presence of *Abyssoanthus* sp. (Anthozoa: Zoantharia) in the Mediterranean Sea: an indication of non-dependence of *Abyssoanthus* to chemosynthetic-based ecosystems. Cahiers De Biologie Marine 51, 475–478.

Springer, M.S., Gatesy, J., 2016. The gene tree delusion. Molecular Phylogenetics and Evolution 94, 1–33.

Stamatakis, A., 2014. RAxML Version 8: A tool for Phylogenetic Analysis and Post-Analysis of Large Phylogenies. Bioinformatics 30, 1312–1313.

Stampar, S.N., Maronna, M.M., Vermeij, M.J.A., Silveira, F.L.D., Morandini, A.C., 2012. Evolutionary diversification of banded tube-dwelling anemones (Cnidaria; Ceriantharia; *Isarachnanthus*) in the Atlantic Ocean. PLoS One 9, e109481.

Streicher, J.W., Schulte, J.A., Wiens, J.J., 2016. How should genes and taxa be sampled for phylogenomic analyses with missing data? An empirical study in iguanian lizards. Systematic Biology 65, 128–145.

Swain, T.D., 2009. Phylogeny-based species delimitations and the evolution of host associations in symbiotic zoanthids (Anthozoa, Zoanthidea) of the wider Caribbean region. Zoological Journal of the Linnean Society 156, 223–238.

Swain, T.D., 2010. Evolutionary transitions in symbioses: dramatic reductions in bathymetric and geographic ranges of Zoanthidea coincide with loss of symbioses with invertebrates. Molecular Ecology 19, 2587–2598.

Swain, T.D., Schellinger, J.L., Strimaitis, A.M., Reuter, K.E., 2015. Evolution of anthozoan polyp retraction mechanisms: convergent functional morphology and evolutionary allometry of the marginal musculature in order Zoanthidea (Cnidaria: Anthozoa: Hexacorallia). BMC Evolutionary Biology 15, 123.

Swain, T.D., Strimatis, A.M., Reuter, K.E., Boudreau, W., 2016. Towards integrative systematics of Anthozoa (Cnidaria): Evolution of form in the order Zoanthidea. Zoologica Scripta DOI:10.1111/zsc.12195.

Swain, T.D., Swain, L.M., 2014. Molecular parataxonomy as taxon description: examples from recently named Zoanthidea (Cnidaria: Anthozoa) with revision based on serial histology of microanatomy. Zootaxa 3796, 81–107.

Swain, T.D., Taylor, D.J., 2003. Structural rRNA characters support monophyly of raptorial limbs and paraphyly of limb specialization in water fleas. Proceedings of the Royal Society B-Biological Sciences 270, 887–896.

Szymanski, M., Erdmann, V.A., Barciszewski, J., 2007. Noncoding RNAs database (ncRNAdb). Nucleic Acids Research 35, D162–D164.

Torres-Suarez, O.L., 2014. *Gorgonia mariae* and *Antillogorgia bipinnata* populations inferred from compensatory base change analysis of the internal transcribed spacer 2. Molecular Phylogenetics and Evolution 79, 240–248.

Wei, N.W.V., Wallace, C.C., Dai, C.F., Pillay, K.R.M., Chen, C.A., 2006. Analyses of the ribosomal internal transcribed spacers (ITS) and the 5.8S gene indicate that extremely high rDNA heterogeneity is a unique feature in the scleractinian coral genus *Acropora* (Scleractinia; Acroporidae). Zoological Studies 45, 404–418.

